# Exploring the Role of MMP9 and its Association with TGFβ Signalling in Patients with Mesial Temporal Lobe Epilepsy-Hippocampal Sclerosis

**DOI:** 10.1101/2023.07.22.550041

**Authors:** Debasmita Paul, Arpna Srivastava, Jyotirmoy Banerjee, Manjari Tripathi, Ramesh Doddamani, Sanjeev Lalwani, Fouzia Siraj, Meher Chand Sharma, Poodipedi Sarat Chandra, Aparna Banerjee Dixit

**Author notes:** Corresponding author: Dr. Aparna Banerjee Dixit, Assistant Professor, Dr. B.R. Ambedkar Center for Biomedical Research, University of Delhi, New Delhi, India Email ID; ORCID ID: http://orcid.org/0000-0003-3028-2259.

## Abstract

Inflammation and blood brain barrier (BBB) damage are associated with epileptogenesis in Mesial Temporal lobe epilepsy with Hippocampal sclerosis (MTLE-HS). Animal studies have predicted the role of Matrix metalloproteinase 9 (MMP9) in extracellular matrix (ECM) modulation, BBB leakage and neuro-inflammation, while Transforming growth factor beta (TGFβ) signalling in astrocytes potentiates hyper-excitability leading to seizure generation. We hypothesize whether changes in activity and expression of MMP9, and the ratio of MMP9 and its inhibitor, Tissue inhibitor of metalloproteinase 1 (TIMP1), have a role in epileptogenesis in the patients with MTLE-HS through zona occludens 1 (ZO1) modulation. We also proposed the role of astrocytic TGFβ signalling in these patients. mRNA expression of MMP9 and TIMP1 was significantly up-regulated. The ratio of MMP9 to its inhibitor TIMP1 was greater than one, suggesting activation of MMP9, further confirmed by gelatin zymography. MMP9 activity as well as immunoreactivity was higher in patients with MTLE-HS as compared to non-seizure controls, whereas the immunoreactivity of ZO1 was significantly lower in the patients. The downstream TGFβ signalling effector molecules, SMAD3 and pSMAD3 immunoreactivity were also significantly higher in MTLE-HS patients and both molecules showed co-localisation with astrocytes in the hippocampal region. Further, we showed preliminary data about interaction of MMP9 and TGFβ1 in these patients as evidenced by a co-immunoprecipitation assay. This study highlighted the MMP9 and astrocytic TGFβ signalling mediated potential mechanism of epileptogenesis in MTLE-HS patients.

## Introduction

Mesial temporal lobe epilepsy with hippocampal sclerosis (MTLE-HS) is one of the most common focal drug resistant epilepsy (DRE). Inflammation and blood brain barrier (BBB) damage have been integral part of the epileptogenesis process in MTLE (Pitkänen and Sutula, 2002). Evidences suggested various roles of matrix metalloproteinases (MMPs) in microglial activation, neuro-inflammation, and production of pro-inflammatory cytokines and BBB leakage (Rempe et al., 2016). Among all the MMPs, MMP9, MMP2 and MMP3 are most abundantly expressed in the central nervous system (Rempe et al., 2016). Recent studies suggest that MMP9 plays an important role in BBB damage and neuro-inflammation thereby regulating synaptic plasticity, learning and memory (Szklarczyk et al., 2002; Nagy et al., 2006; Tian et al., 2007).

MMP9 is found in a variety of cell types throughout the body, including neurons, glia, and endothelial cells (Genersch et al., 2000). It forms a complex with tissue inhibitor of matrix metalloproteinase (TIMP1), an endogenous inhibitor of MMP9, prior to secretion, and is secreted at levels comparable to MMP9 (Roderfeld et al., 2007). MMP9 gets activated from its pro-form by a cascade involving stromelysin and MMP3 (Suenaga et al., 2000) and this tight regulation is responsible for extracellular matrix (ECM) remodelling that lead to a variety of neurological disorders, including epilepsy. The aberrant plasticity may contribute to neuronal loss in hippocampi, astrogliosis, aberrant pruning of dentate gyrus. ECM is involved in the formation and maintenance of stable neural network connections that guide synaptic plasticity, whereas its destruction, particularly by MMP9, results in abnormal connections us (DG) spines (Wilczynski et al., 2008), reorganization of interneuron terminals (Pollock et al., 2014), loss of perineuronal net (PNN) structures (Pollock et al., 2014, McRae et al., 2012), and the sprouting of mossy fibres that all eventually turn to form a recurrent network (Wilczynski et al., 2008). BBB leakage was linked to higher MMP9 protein and activity levels in the blood and CSF of patients with generalised tonic–clonic seizures (Li et al., 2013). MMP9 may cause BBB degradation by degrading tight junctional (TJ) proteins, zona occludens 1 (ZO1) and occludin (Asahi et al., 2001). These studies suggest MMP9 might play a critical role in epileptogenesis and seizure generation.

An earlier study highlighted activation of TGFβ signalling in patients with MTLE-HS (Paul et al., 2018). Increased expression of TGFβ1 ligand, TGFβRII, SMAD3 and pSMAD3 were demonstrated in patients with MTLE-HS (Paul et al., 2018). BBB dysfunction cause changes in the expression of astrocytic genes such as ion channels and water channels, as well as up-regulation of pro-inflammatory cytokines mediated by TGFβ signalling (Cacheaux et al., 2009) and excitatory synaptogenesis in astrocytic cultures (Weissberg et al., 2015), all of which promote epileptogenesis (Weissberg et al., 2015). The ECM modifying molecules MMP9, MMP14, and TIMP1 were reported to be up-regulated as a result of albumin mediated BBB opening. TGFβ, like albumin, causes BBB disruption in animal models, which could lead to conjecture that TGFβ interacts with MMP9 (Cacheaux et al., 2009). Taking all of this into account, this study was aimed to investigate MMP9 expression, its activity, and localization in neurons and glia, which may contribute to epileptogenesis in MTLE-HS patients via ZO1 modulation. Ratio of MMP9 to its inhibitor TIMP1 was also determined. The interaction of MMP9 and TGFβ1 was also investigated. We also studied the localization of the downstream molecules of TGFβ signalling, SMAD3 and pSMAD3, in astrocytes in patients with MTLE-HS due to a dearth of data on astrocytic TGFβ signalling in brain tissue samples of DRE patients.

## Methods

### Patients and controls

The patients (31 males and 10 females) who were diagnosed to have DRE due to MTLE-HS and underwent resective surgery from January 2015 to December 2019 were included in the study. Each patient had a pre-surgical evaluation, and the pathology was determined by analysing convergent data from MRI, video electroencephalogram (vEEG), Fluoro-2- deoxyglucose positron emission tomography (FDG-PET), and magnetoencephalography (MEG), as well as histopathological examinations by a neuropathologist. Patients were included in the study if they had MRI-vEEG-MEG concordance, no visible structural changes/pathologies in extra-hippocampal areas, and no dual pathology (e.g. cortical dysplasia) (Tripathi et al., 2016; Paul et al., 2018). We have used histologically normal hippocampal tissues obtained from post-mortem cases without any history of seizures or other neurological disorders as non-seizure controls. All the autopsies were performed within 12 h of death (Paul et al., 2018). A part of the resected tissue was preserved in 4% paraformaldehyde (PFA) for immunostaining, while the remainder was frozen and stored at −80°C until use. Table 1 shows the clinical features of the patients and controls.

**Table 1:** Clinical characteristics of patients with MTLE-HS and non-seizure controls.

| Patient | Age | Sex | Pathology | Age at onset of seizure (years) | Anti epileptic drugs (AED) | Engel surgical outcome scale (Engel, 1995) |
| --- | --- | --- | --- | --- | --- | --- |
| M1 | 11 | M | MTLE-HS | 1.5 | PHE,OXC,LEV,CLO | II |
| M2 | 4 | M | MTLE-HS | 0.6 | LTG,VAL,CLO | III |
| M3 | 13 | M | MTLE-HS | 5 | CLO, LTG | I |
| M4 | 22 | M | MTLE-HS | 12 | CLO, LTG,VAL | I |
| M5 | 51 | F | MTLE-HS | 18 | CLO,LEV | I |
| M6 | 22 | M | MTLE-HS | 19 | LEV, CLO | I |
| M7 | 25 | M | MTLE-HS | 15 | LEV, VAL | I |
| M8 | 16 | M | MTLE-HS | 6 | CLO, PHE, CAR | I |
| M9 | 29 | M | MTLE-HS | 19 | VAL, CAR | I |
| M10 | 19 | F | MTLE-HS | 12 | CLO, OXC, LEV | I |
| M11 | 30 | M | MTLE-HS | 16 | LEV, CAR | I |
| M12 | 46 | M | MTLE-HS | 23 | LEV, OXC, CLO | I |
| M13 | 51 | F | MTLE-HS | 46 | CAR, LTG, CLO | I |
| M14 | 35 | M | MTLE-HS | 3 | CAR,CLO,LEV | I |
| M15 | 34 | M | MTLE-HS | 26 | LEV,CAR,CLO | I |
| M16 | 42 | F | MTLE-HS | 28 | LEV, LCS | I |
| M17 | 43 | M | MTLE-HS | 21 | CLO,LEV,CAR | I |
| M18 | 20 | F | MTLE-HS | 12 | CLO,LEV | I |
| M19 | 24 | M | MTLE-HS | 19 | LEV,OXC, LTG | II |
| M20 | 22 | M | MTLE-HS | 15 | CLO,LCS,OLA | I |
| M21 | 36 | M | MTLE-HS | 20 | VAL, LTG | I |
| M22 | 9 | M | MTLE-HS | 1 | PHE,CAR | I |
| M23 | 27 | F | MTLE-HS | 25 | LEV,CLO | I |
| M24 | 6 | M | MTLE-HS | 0.2 | CLO,LEV,PHE,VAL,<br>CLN | I |
| M25 | 22 | M | MTLE-HS | 8 | CLO,LEV | I |
| M26 | 36 | F | MTLE-HS | 19 | OXC,CAR | I |
| M27 | 24 | M | MTLE-HS | 0.6 | CAR,CLO,CLN,VAL,<br>TPR | I |
| M28 | 24 | F | MTLE-HS | 1.5 | PHE,TPR | I |
| M29 | 6 | M | MTLE-HS | 2 | CLO, LTG,VAL | I |
| M30 | 31 | F | MTLE-HS | 20 | CAR,CLO,CLN | I |
| M31 | 21 | M | MTLE-HS | 15 | LEV,CLO | I |
| M32 | 23 | M | MTLE-HS | 3 | OXC, LCS, CLO | I |
| M33 | 39 | M | MTLE-HS | 5 | CAR,LCS,LEV,LTG | I |
| M35 | 18 | M | MTLE-HS | 6 | CAR, VAL,CLO | I |
| M36 | 24 | M | MTLE-HS | 2.5 | CAR,LCS,LEV,LTG | I |
| M37 | 36 | F | MTLE-HS | 5 | LEV,CAR,PHE,TPR | III |
| M38 | 26 | M | MTLE-HS | 2 | CAR, VAL,CLO | I |
| M39 | 52 | M | MTLE-HS | 39 | LEV, PHE | I |
| M40 | 26 | M | MTLE-HS | 2 | CAR, VAL, CLO | I |
| M41 | 21 | M | MTLE-HS | 10 | CAR, CLN | I |
| A1 | 32 | M | ABDOMINAL<br>INJURIES |  | N.A. |  |
| A2 | 20 | M | HANGING |  | N.A. |  |
| A3 | 16 | M | CHRONIC<br>LIVER<br>DISEASE |  | N.A. |  |
| A4 | 26 | M | ASPHYXIA |  | N.A. |  |
| A5 | 26 | F | HANGING |  | N.A. |  |
| A6 | 25 | M | MYOCARDIAL<br>INFARCTION |  | N.A. |  |
| A7 | 19 | M | ABDOMINAL<br>&PELVIC<br>INJURY |  | N.A. |  |
| A8 | 19 | M | ASPHYXIA |  | N.A. |  |
| A9 | 18 | M | POISONING |  | N.A. |  |
| A10 | 28 | M | ASPHYXIA |  | N.A. |  |
| A11 | 48 | M | MYOCARDIAL<br>INFARCTION |  | N.A. |  |
| A12 | 1.6 | M | DROWNING |  | N.A. |  |
| A13 | 65 | F | MYOCARDIAL<br>INFARCTION |  | N.A. |  |
| A14 | 18 | F | HANGING |  | N.A. |  |
| A15 | 41 | M | ABDOMINAL INJURY |  | N.A. |  |
| A16 | 20 | M | HANGING |  | N.A. |  |
| A17 | 42 | M | PELVIC, LIMB INJURY |  | N.A. |  |
| A18 | 6 | M | HANGING |  | N.A. |  |
| A19 | 24 | M | HANGING |  | N.A. |  |
| A20 | 14 | M | ELECTROCUTION |  | N.A. |  |
| A21 | 51 | M | SUDDEN DEATH |  | N.A. |  |
| A22 | 35 | M | MYOCARDIAL INFARCTION |  | N.A. |  |
| A23 | 17 | M | DROWNING |  | N.A. |  |

M=MTLE-HS, A= Autopsy Control, COD= Cause of death, CLO= clobazam, VAL= Valproate, LEV= Leviteracetum, OXC= Oxcarbamazepine, CAR= Carbamazepine, PHE= Phenytoin, LTG=Lamotrigen, PBT= Phenobarbital, CLN= Clonazepam, LCS= Lacosamide, TPR= Topiramate

### Hematoxylin and eosin (H&E) staining

The procedures were conducted as described previously (Canene-Adams, 2013; Zhou et al., 2016). The resected tissue samples from MTLE-HS and autopsy hippocampal samples were fixed in 4% paraformaldehyde. After being embedded in paraffin and cut into 5μm sections, slides were stained with H&E to observe the histological changes. The sections were examined using an OLYMPUS BX53 Upright Research Microscope.

### Quantitative real-time polymerase chain reaction (RT-qPCR)

RT-qPCR was performed to evaluate the mRNA expression level of MMP9 and TIMP1. Total RNA was isolated using RNeasy Mini kit (Qiagen) as per manufacturer’s protocol. RNA was estimated using Qubit 3.0 fuorometer (Life Technologies). cDNA synthesis was done using high-capacity cDNA synthesis kit according to the manufacturer’s recommendations (ThermofischerScientifc). HPRT (hypoxanthine phosphoribosyl-transferase) was included as reference gene. Specific primers were designed using the Primer-BLAST (Primer3Input, version 0.4.0 and BLAST, available at http://www.ncbi.nlm.nih.gov/tools/primer-blast/). Following are the sequences: [MMP9-F-5’CGTCTTCCCCTTCACTTTCCTG-3’];[MMP9-R- 5’CCCCACTTCTTGTCGCTGTC-3’];TIMP1-F-5’TCCCAGATAGCCTGAATCCT-3’];[TIMP1-R 5’CTGCTGGGTGGTAACTCTTTAT-3’]. The template was amplified at 95°C for 3 min, followed by 40 cycles of 95 °C for 5 sec and 60 °C for 30 sec using the CFX 96 real-time systems (Biorad). The 2−ΔΔCt method was used to analyze the data (Schmittgen and Livak, 2008).The melting curve of each replicate was examined to confirm a single peak appearance.

### Enzyme linked immunosorbent assay (ELISA)

Enzyme-linked immunosorbent assay (R&D System, United States) was performed according to the instructions to measure the protein expression level of MMP9 and TIMP1 in resected hippocampi of non-seizure controls and patients with MTLE-HS. 200μg protein was added into pre-coated plates for overnight incubation at 4ᵒC, followed by incubation for 2 h at 37ᵒC with biotinylated secondary antibody. The absorbance of each well at 562nm was measured and the ratio of MMP9 and TIMP1 was calculated. Value was expressed as nanograms of MMP9/TIMP1 per milligram (ng/mg) protein.

### Gelatin Zymography

Zymography was performed essentially as described previously (Frankowski et al., 2012). Equal amount of protein (25μg) from each sample was subjected to 10% SDS-PAGE gel containing 0.5% gelatin under non-denaturing, non-reducing conditions. Following electrophoresis, gels were washed in renaturation buffer (2.5% Triton X-100 in 50 mm Tris– HCl (pH 7.5)) for 1 h in an orbital shaker. Then the zymograms were incubated for 18 h at 37°C in incubation buffer (0.15 m NaCl, 10 mm CaCl2, 0.02% NaN3 in 50 mm Tris–HCl (pH 7.5). Gels were then stained with Coomassie blue and destained with 7% methanol and 5% acetic acid. Enzymatic activity was visible as clear bands against a dark background.

### Immunohistochemistry and Double-Immunofluorescence

Immunohistochemical staining (IHC) was done using the ABC Elite Vectastain kit (Vectorlab, PK6101) following the manufacture’s protocol. Briefly, sections were first deparaffinized with absolute xylene, followed by rehydration with absolute ethanol in a gradient series. Antigen retrieval was performed in 0.1% citrate buffer for 30 min and then sections were quenched in hydrogen dioxide (3%) solution for 15 min. Blocking was done using normal goat serum (3%– 5%) containing 0.1% Triton X-100. Primary antibodies, human anti-SMAD3 (1:200) (Abcam, ab40854), anti-pSMAD3 (1:150) (Abcam, ab52903), anti-MMP9 (1:100) (Abcam, ab38898), human anti-ZO1 (1:300), (Thermo Fischer, 61-7300) were incubated overnight at 4°C, and further incubated for 90 min with the appropriate secondary antibody. Following a 0.1M PBS rinse, the slides were stained with a DAB solution and dehydrated with a graded ethanol series. The sections were mounted with DPX and observed under the bright field of the BX53 Upright Research Microscope (OLYMPUS). Image J software was used to quantitatively analyse the positive expression of each protein detected by immunohistochemistry. The average number of cells expressing the protein of interest in patients with MTLE-HS and controls was calculated by counting ten fields from two different sections on the same slide from the same sample.

For double immuno-labelling, the cell membranes were permeabilised with 0.3% Triton X-100 for 15 min. Blocking was performed with 5% goat serum for 2 h at 25°C. Sections were incubated with primary antibodies at 4°C for overnight. Primary antibodies used were: (human anti-SMAD3 (1:1:200) (Abcam, ab40854), and anti-pSMAD3 (1:150) (Abcam, ab52903), human anti-GFAP (1:400) (Abcam, ab53554)). The sections were incubated for 2 h at 25°C with appropriate secondary florescence labelled antibodies (donkey anti-goat IgG, Alexa fluor 488 (Abcam, ab150129) & donkey anti-rabbit IgG, Alexa fluor 594, (Abcam, ab150076) (1:800) for 2 h in the dark at RT and counterstained with DAPI (1:1000, Sigma, T7024). Finally, the sections were observed with a fluorescence microscope BX53 Upright Research Microscope (OLYMPUS).

### Co-immunoprecipitation

For analysing protein-protein interaction, we performed the co-immunoprecipitation experiments (Zhou et al., 2016) using hippocampal samples from non-seizure controls and patients with MTLE-HS. The proteins were extracted in RIPA buffer and estimated using Bicinchoninic acid (BCA) kit. 300μg of protein was used for immunoprecipitation assay using immunoprecipitation kit containing Dynabeads-coupled protein G (magnetic beads) and human anti-MMP9 (1:700) (Abcam, ab38898). Briefly the protein was mixed with anti-MMP9 antibody and incubated overnight at 4°C, then 50μl beads were added in the solution. The pellet obtained was washed with 0.01M PBS containing 0.1% Tween 20 and run on 10% SDS-PAGE under denaturing conditions. Then western blot was performed using the primary antibody against human anti-TGFβ1 (1:500) (Abcam, ab122975) as described previously (Paul et al., 2018). For positive control, 100μg of total protein was used in the western blot.

### Statistical analysis

Student’s t-test was used to determine statistical significance for mRNA expression of MMP9 and TIMP1, gelatinase activity of MMP9 and ratio of MMP9:TIMP1, IHC for GFAP, NeuN, SMAD3, MMP9 and ZO1, while protein expression of MMP9 and TIMP1 and IHC for pSMAD3 was analysed by Mann Witney U test using SigmaPlot13.1software (Systat Software, Inc., Chicago, IL). TGFβ1 and pSMAD3 expression were correlated with epilepsy duration using Spearman correlation (R). The data was presented as a mean with standard deviation. A p-value of <0.05 was considered statistically significant.

## Results

### Histological findings of non-seizure controls and patients with MTLE-HS

We had observed no loss of neurons and gliosis (Figure 1A, a) through H&E staining of hippocampal sections of non-seizure controls while H&E staining of hippocampal sections of patients with MTLE-HS showed neuronal loss (Figure 1A, b), gliosis (Figure 1A, c) and edema (Figure 1A, d). The hippocampal sections of MTLE-HS showed prominent astrogliosis (Figure 1B, c). The immunohistochemical staining of the hippocampal sections of non-seizure controls showed morphologically normal astrocytes and significantly lower GFAP IR (12.25±0.739087) (Figure 1B, a) as compared to MTLE-HS patients (19.59583±0.956088, p=<0.001) (Figure 1B, b). The Hippocampal sections of MTLE-HS are characterized by astrogliosis and hypertrophy of primary processes, and an increase in the expression of GFAP (Figure 1B, b). The immunohistochemical staining of the hippocampal sections of non-seizure controls showed neurons distributed throughout the field with significantly high NeuN IR (10.78333±0.433205) (Figure 1B, d) compared to hippocampal sections of MTLE-HS (4.775±0.48657, p=<0.001) (Figure 1B, e). Marked loss of neurons was seen in hippocampal sections of MTLE-HS patients (Figure 1B, e).

**Figure 1:**
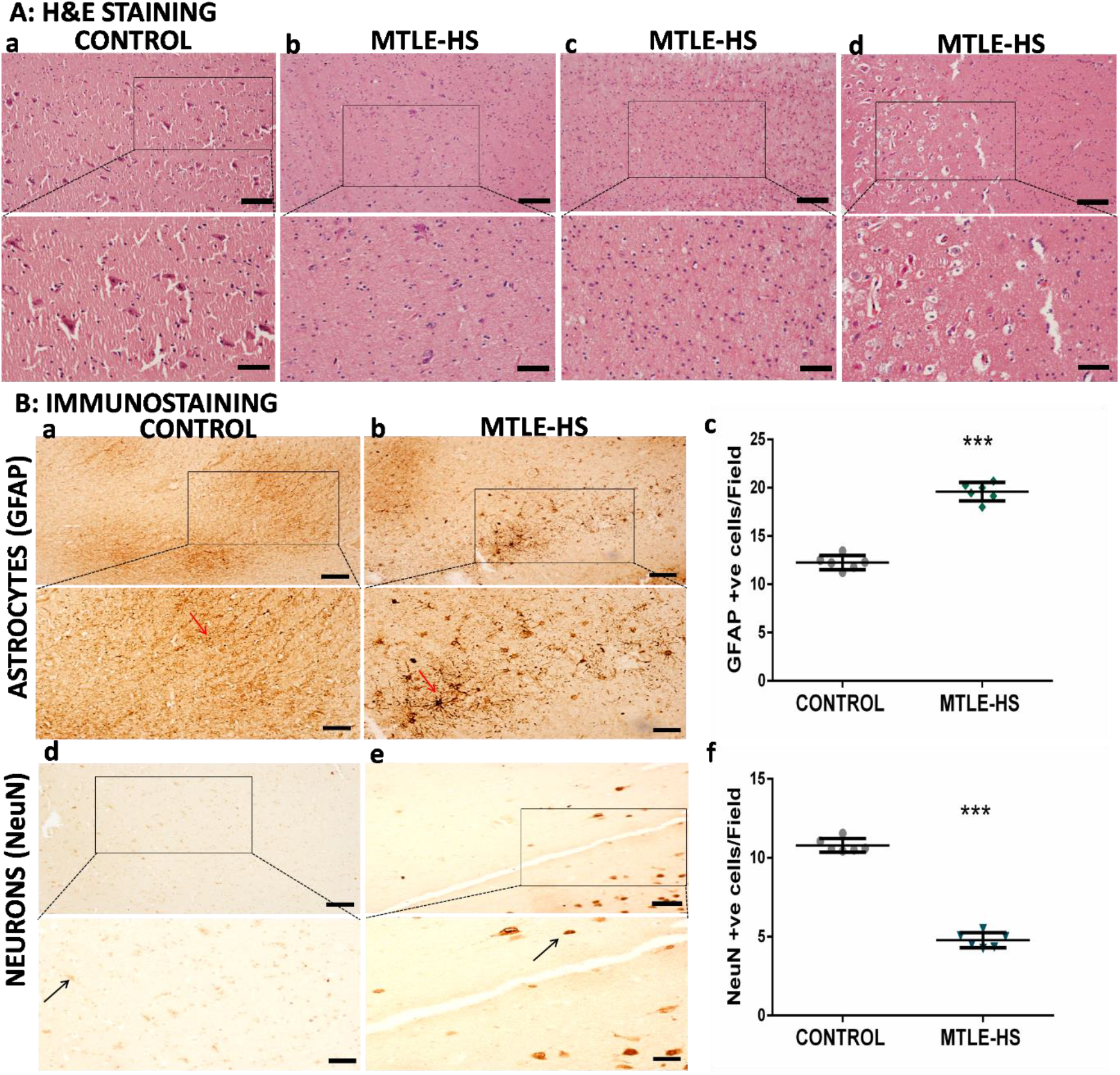
Representative photomicrographs of H&E staining and immunostaining of hippocampal tissue of controls and patients with MTLE-HS. A) H&E staining of the hippocampal sections of non-seizure controls (1A-a) and patients with MTLE-HS (1A-b, c, d). B) Upper panel-Immunostaining of the hippocampal sections of non-seizure Controls (1B-a, n=6) and patients with MTLE-HS (1B-b, n=6) using GFAP as marker for astrocytes; Lower panel-Immunostaining of the hippocampal sections of non-seizure Controls (1B-d, n=6) and patients with MTLE-HS (1B-e, n=6) using NeuN as marker for neurons. The graphical representation of GFAP+ cells (1B-c) and NeuN+ cells (1B-f) were shown by scatter plots in controls and patients. Black arrows showed neurons and red arrows show astrocytes. The boxed area of the images (10X) had been magnified (20X) and connected through dotted lines. Error bar represented mean ± SD. Student’s t-test was used for analysis of NeuN and GFAP IR (p=<0.001). Scale bar, 100μm (scale bar in inset, 50μm)

### Alteration of gene expression and ratio of MMP9 and TIMP1, and activity of MMP9 in patients with MTLE-HS

MMP9 mRNA expression (4.75fold; ranges from 0.74 fold to 26.72 fold, p=0.044) and TIMP1 mRNA expression (3.15 fold; ranges from 0.83 fold to 7.67 fold, p=0.031) were significantly higher in MTLE-HS as compared to non-seizure controls (Figure 2A).

**Figure 2:**
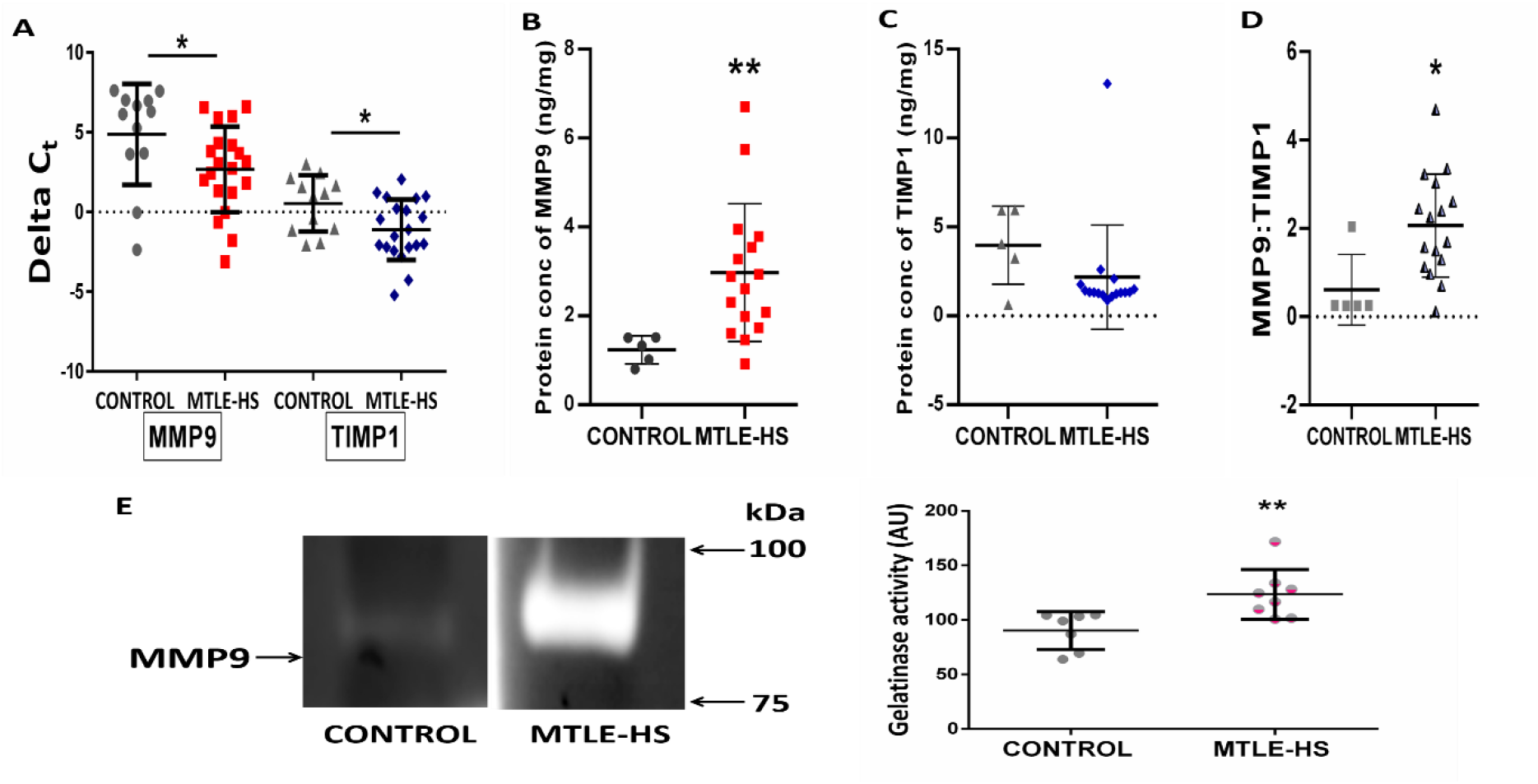
Altered Transcript and protein expression of MMP9 and TIMP1 and its ratio, enzymatic activity of MMP9 in patients with MTLE-HS. A) MMP9 and TIMP1 mRNA expression were analysed by quantitative reverse transcriptase Real Time PCR in hippocampi of patients with MTLE-HS (n=20) as compared to hippocampi of non-seizure controls (n=12). Scatter plot represents data as delta cycle threshold (ΔCt) values and each sample was analysed in triplicates. B&C) MMP9 and TIMP1 protein expression were analysed by ELISA in hippocampi of patients with MTLE-HS (n=16) as compared to hippocampi of non-seizure controls (n=5). Scatter plot represented the distribution of protein expression of MMP9 and TIMP1 and each sample was analysed in triplicates. D) Scatter plot represented the ratio of MMP9:TIMP1 in hippocampi of non-seizure controls and patients with MTLE-HS. E) Zymogram represented enzymatically active MMP9 (∼ 82kDa) from the pro MMP9 form, which is visualised as bright bands against dark background Graphical representation of activity of MMP9 in the resected hippocampi of patients with MTLE-HS (n=8) as compared to hippocampi of non-seizure controls (n=7). The images were adjusted to an external grey scale. Error bar represented mean ± SD. Student’s t-test was performed for MMP9 mRNA transcripts (p=0.044), TIMP1 mRNA transcripts (p=0.031), MMP9:TIMP1 ratio (p=0.0183) and MMP9 activity (p=0.00813). Mann Witney U test was performed for MMP9 (p=0.006), TIMP1 (p=0.107) protein expression.

Progress of activation of pro-MMP9 was in part determined by the ratio of MMP9 to TIMP1. When the MMP9/TIMP1 ratio exceeds 1.0, MMP9 activation might happen by plasmin/stromelysin1 protease cascade (Ramos et al., 1999). We analysed the protein expression by ELISA and found that MMP9 expression was significantly higher in MTLE-HS (2.297±1.54 ng/mg, p=0.006) vs autopsy controls (1.234±0.31 ng/mg) (Figure 2B) while TIMP1 protein expression (2.174±2.93ng/mg, p=0.107) was unaltered (Figure 2C). The ratio of MMP9 and TIMP1 exceeded 1 (2.06±0.527, p=0.0183) in the hippocampi of patients with MTLE-HS indicating probable activation of MMP9 in these patients (Figure 2D). MMP9 is known to degrade gelatin thereby its activity was accessed by Gelatin zymography where the MMP9 activity was detected as bright bands against dark background. These bright bands appeared in areas where active MMP9 lead to degradation of gelatin (Figure 2E). We have also detected a band corresponding to 150 kDa and 220kDa (data not shown) and might represent MMP9 dimer or complex formed with other proteins (Woessner et al., 1995). The activity of MMP9 was significantly higher in patients with MTLE-HS (123.35±22.95, p=0.00813) as compared to non-seizure controls (90.208±17.28) (Figure 2E).

### Higher SMAD3 and pSMAD3 Immunoreactivity (IR) and co-localisation of SMAD3/pSMAD3 with the astrocytes implies role of astrocytic TGFβ signalling in patients with MTLE-HS

Based on study portraying higher protein expression of SMAD3 and its phosphorylated form, pSMAD3 in the total protein lysate in patients with MTLE-HS (Paul et al., 2018), we measured the expression of SMAD3 and pSMAD3 in different cells of the resected hippocampi by immunological staining. We observed that SMAD3 IR was significantly higher in the hippocampi of patients with MTLE-HS (7.191667±1.742819, p=<0.001) as compared to the hippocampi of non-seizure controls (2.058333±0.451017) (Figure 3A-C). Also, pSMAD3 IR was significantly higher in the hippocampi of patients with MTLE-HS (20.76±3.92349, p=<0.001) as compared to the hippocampi of controls (6.2±1.3775). pSMAD3 showed nuclear localisation in all the cells (Figure 4A-C). Consequently, we investigated whether TGFβ signalling was linked to astrocytes. We performed the double immunofluorescence study using GFAP as a marker for the reactive astrocytes, and SMAD3 & pSMAD3 as downstream effector of TGFβ signalling. Our results demonstrated that GFAP co-localises with both SMAD3 (Figure 5A) and pSMAD3 (Figure 5B) in the hippocampus of the patients with MTLE-HS. However, in the Control hippocampus, absence of co-labelling of GFAP with SMAD3 (Figure 5A) and pSMAD3 (Figure 5B) were observed. This co-expression of SMAD3 and pSMAD3 with astrocytes suggested for the activation of astrocytic TGFβ signalling in patients with MTLE-HS.

**Figure 3:**
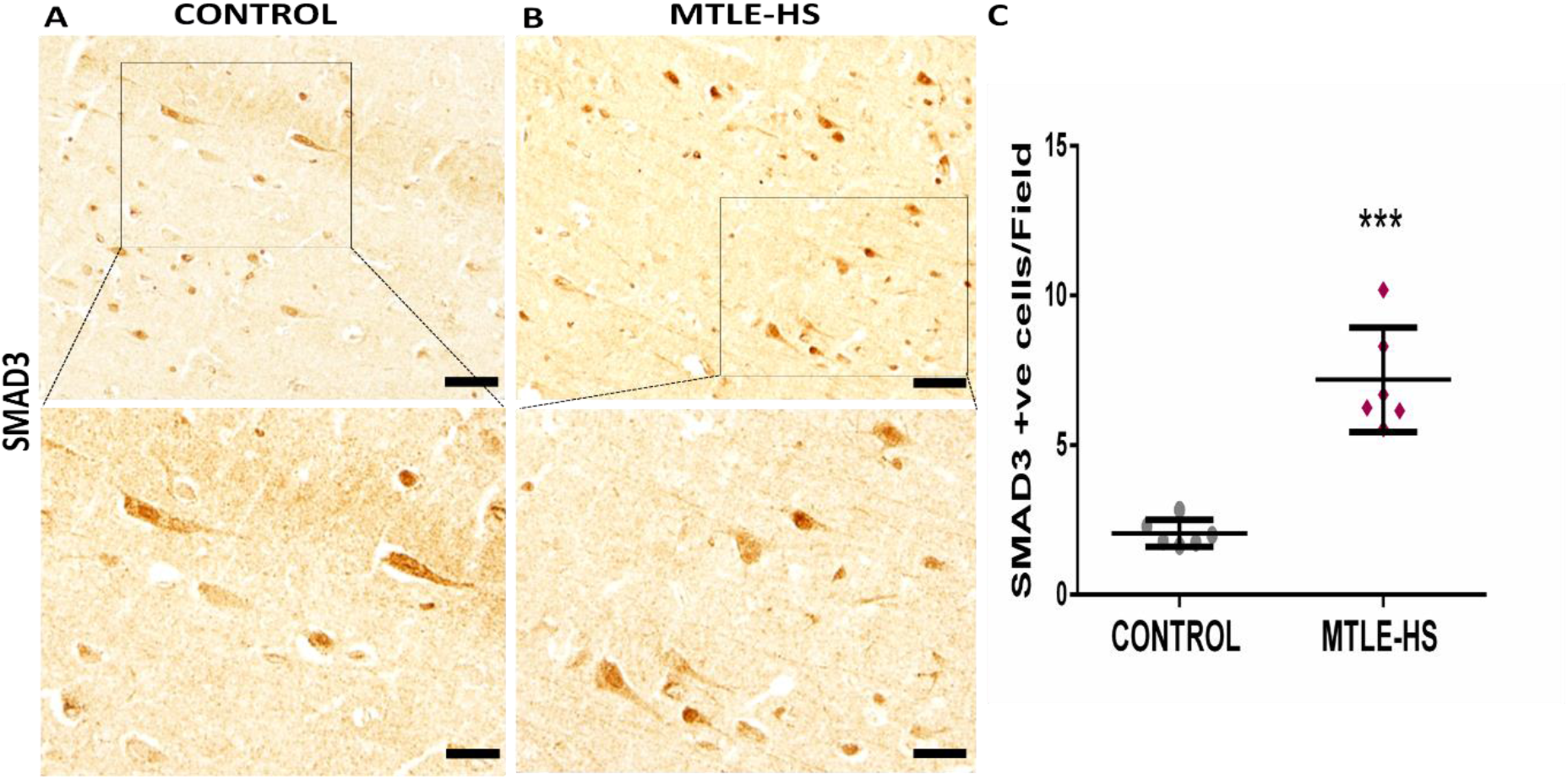
Higher SMAD3 immunoreactivity in resected hippocampi of patients with MTLE-HS. Representative photomicrographs of immunostaining of A) hippocampal sections of non-seizure controls (n=6) and B) hippocampal sections of patients with MTLE-HS (n=6). C) Scatter plot depicted number of cells positively expressing SMAD3 IR in controls and patients with MTLE-HS. The boxed area of the images (20X) had been magnified (40X) and connected through dotted lines. Error bar represented mean ± SD. Student’s t-test was applied to test the statistical significance for SMAD3 IR (p<0.001). Scale bar 50μm (scale bar in inset, 20μm)

**Figure 4:**
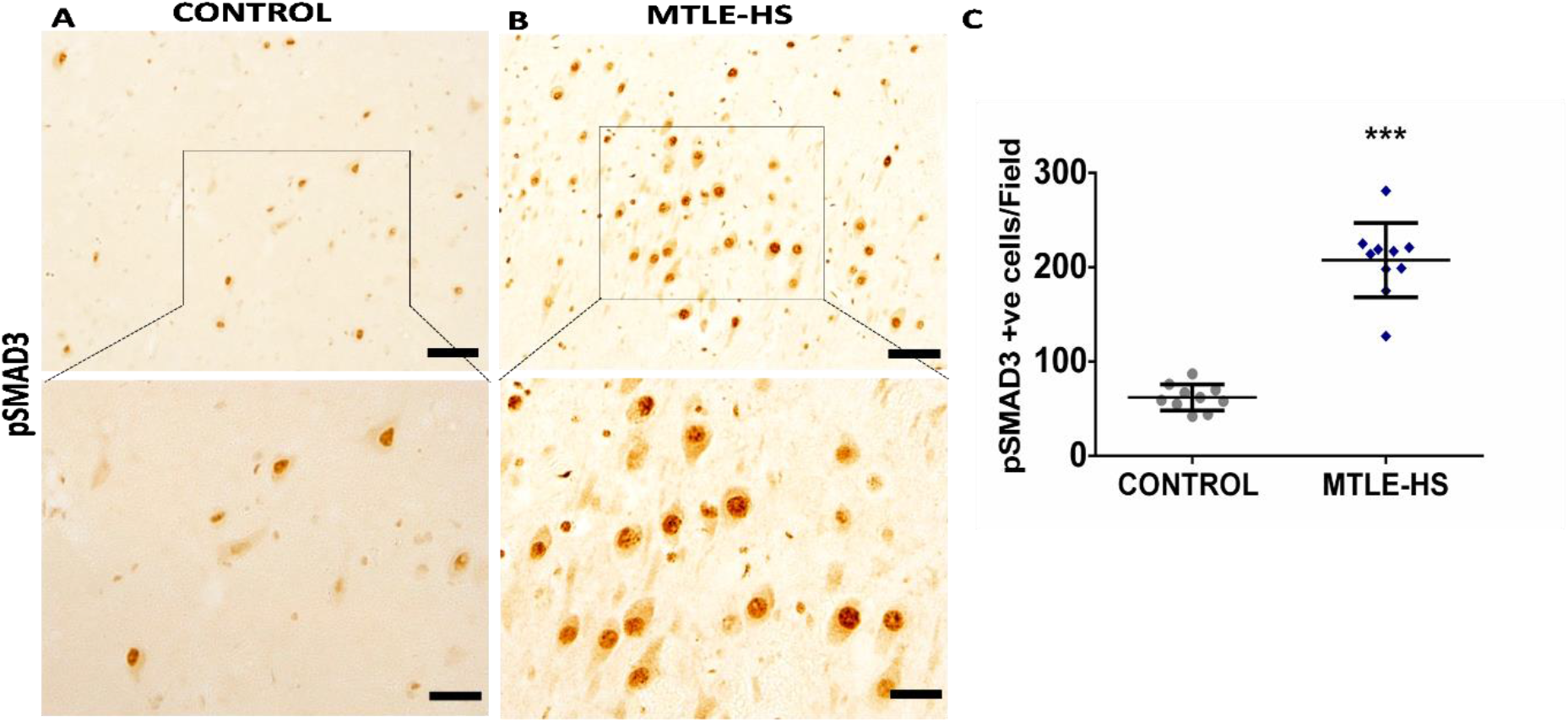
Higher pSMAD3 immunoreactivity in resected hippocampi of patients with MTLE-HS. Representative photomicrographs of immunostaining of A) hippocampal sections of non-seizure controls (n=10) and B) hippocampal sections of patients with MTLE-HS (n=10). C) Scatter plot depicted number of cells positively expressing pSMAD3 IR in controls and patients with MTLE-HS. The expression of pSMAD3 is nuclear as shown in (A&B). The boxed area of the images (20X) had been magnified (40X) and connected through dotted lines. Error bar represented mean ± SD. Mann Witney U test was used to analyze the pSMAD3 IR (p=<0.001). Scale bar 50μm (scale bar in inset, 20μm)

**Figure 5:**
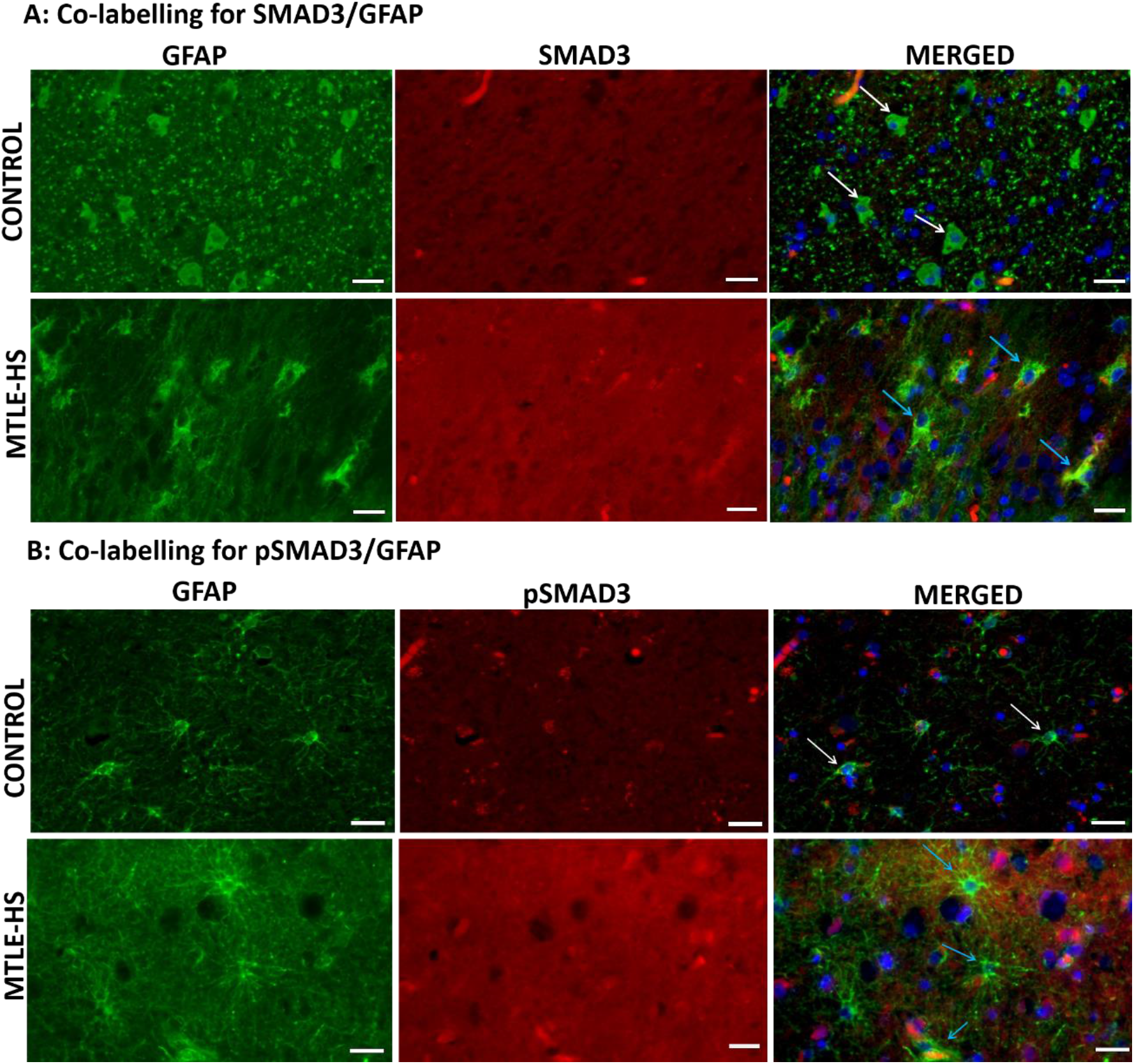
Double immunostaining depicts co-labelling of GFAP along with SMAD3 and pSMAD3 in resected hippocampi of patients with MTLE-HS. Representative photomicrographs of co-labelling of GFAP (an astrocyte marker) with A) SMAD3 in the hippocampi of non-seizure controls (n=6) and patients with MTLE-HS (n=6) and with B) pSMAD3 in the hippocampi of non-seizure controls (n=6) and patients with MTLE-HS (n=6). Astrocytes are stained with GFAP (green) and SMAD3/pSMAD3 proteins with anti-SMAD3/pSMAD3 antibodies (red). In upper panel, A) In Controls, no co-localisation of GFAP and SMAD3 (indicated by white arrows) was observed while in patients with MTLE-HS, co-localisation of GFAP and SMAD3 (yellow) was observed (indicated by blue arrows). In lower panel, B) In Controls, no co-localisation of GFAP and pSMAD3 (indicated by white arrows) was observed while in patients with MTLE-HS, co-localisation of GFAP and pSMAD3 (yellow) was observed (indicated by blue arrows). Scale bar, 20μm.

### Higher MMP9 and lower ZO1 immunoreactivity (IR) in patients with MTLE-HS corroborate to BBB damage

The cellular localisation of MMP9 in the hippocampus was determined to confirm that expression and activity of MMP9 was pertinent. We observed that the number of glia and neuronal cells expressing MMP9 IR was significantly higher in the hippocampi of patients with MTLE-HS (23.1625±2.09961, p=<0.001) as compared to the hippocampi of controls (9.2±1.30384) (Figure 6A-C). This further confirmed the high expression of MMP9 in the patients. Tight junctions (TJs) are a key dynamic feature of the BBB, and ZO1 is the main protein whose loss affects BBB permeability. We observed that ZO1 IR was significantly lower in the hippocampi of patients with MTLE-HS (11.12±3.1378, p=0.004) as compared to the hippocampi of controls (16.75±3.0755) (Figure 7A-C). The MMP9 IR was observed in the endothelial cell lining of the micro blood vessels present throughout the hippocampus of patients with MTLE-HS (Supplementary 1-A). The ZO1 IR was observed in the endothelial cell lining of the micro blood vessels present throughout the hippocampus of controls and patients. ZO1 expression was prominent in intact vessels in hippocampus of controls (Supplementary 1-B) while its expression was disrupted in damaged vessels in the sclerotic hippocampus of the patients (Supplementary 1-C).

**Figure 6:**
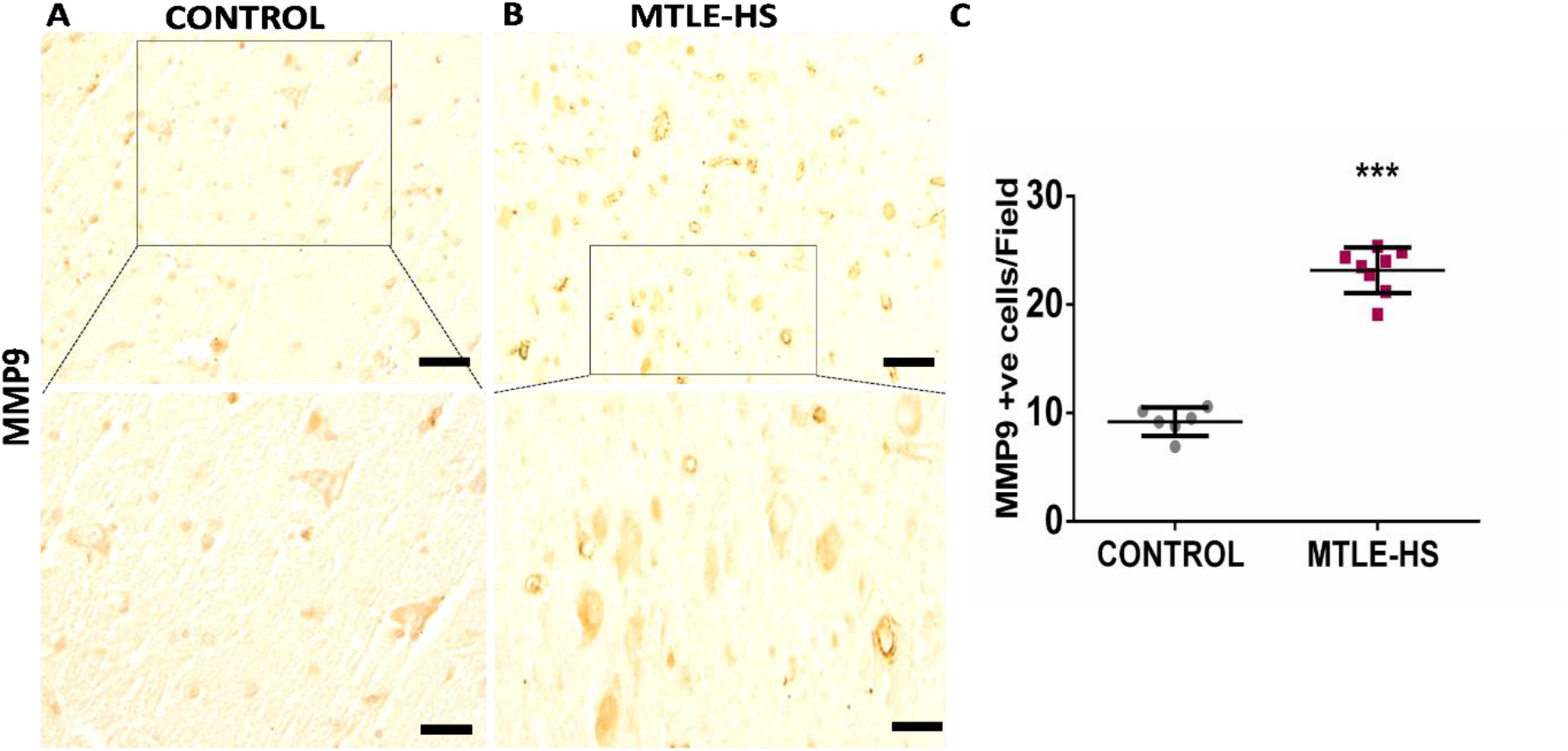
Higher MMP9 immunoreactivity in the resected hippocampi of patients with MTLE-HS. Representative photomicrographs of immunostaining of A) hippocampal sections of non-seizure controls (n=6) and B) hippocampal sections of patients with MTLE-HS (n=8). C) Scatter plot depicted number of cells positively expressing MMP9 IR in controls and patients with MTLE-HS. The boxed area of the images (20X) had been magnified (40X) and connected through dotted lines. Error bar represented mean ± SD. Student’s t-test, p=<0.001 for MMP9 IR. Scale bar 50μm (scale bar in inset, 20μm)

**Figure 7:**
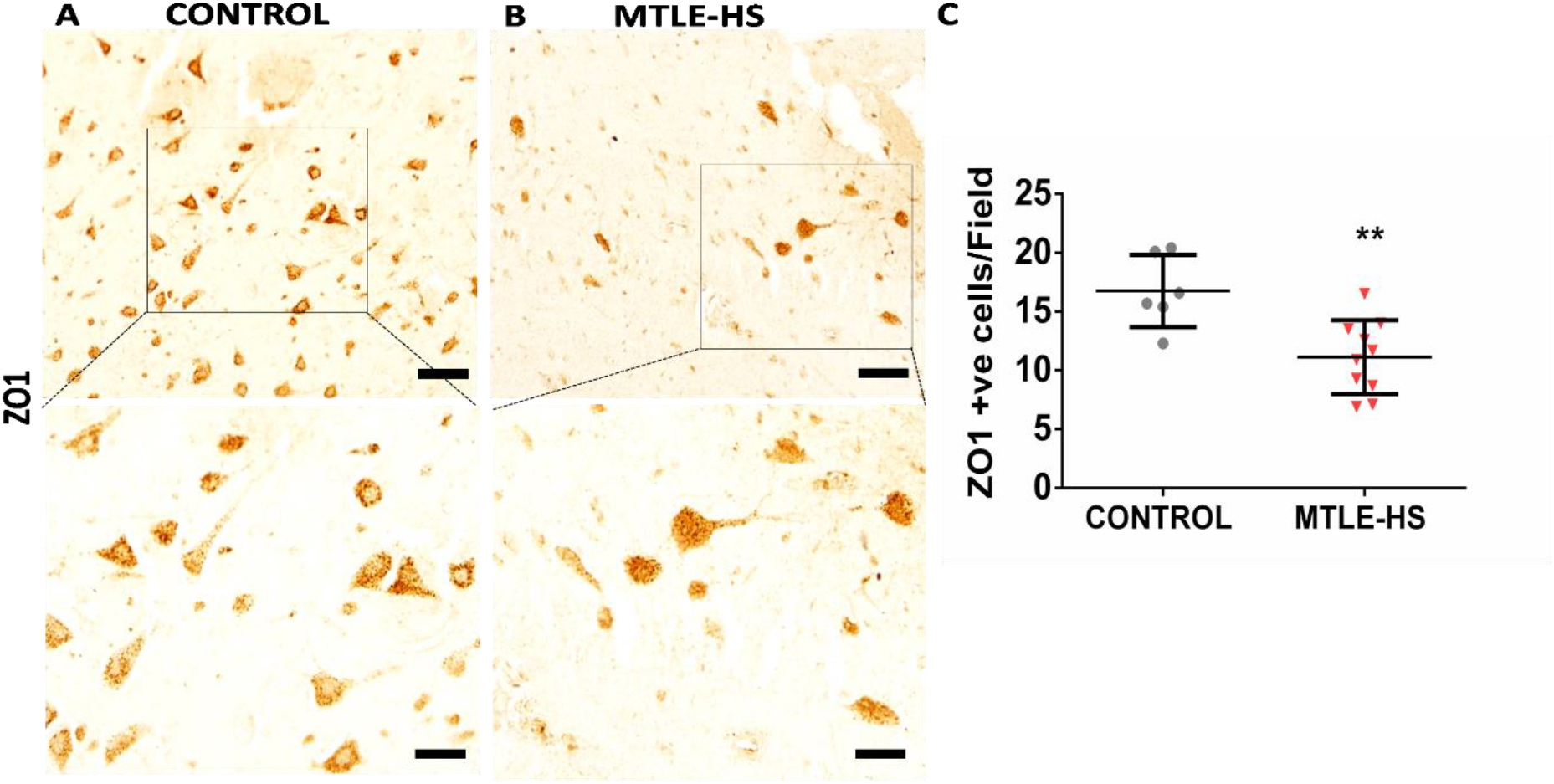
Lower ZO1 immunoreactivity in the resected hippocampi of patients with MTLE-HS. Representative photomicrographs of immunostaining of A) hippocampal sections of non-seizure controls (n=6) and B) hippocampal sections of patients with MTLE-HS (n=10). C) Scatter plot depicted number of cells positively expressing ZO1 IR in controls and patients with MTLE-HS. The boxed area of the images (20X) had been magnified (40X) and connected through dotted lines. Error bar represented mean ± SD. Student’s t-test, p=0.004 for ZO1 IR. Scale bar 50μm (scale bar in inset, 20μm)

### MMP9 and TGFβ1 might be associated in patients with MTLE-HS

Studies show that astrocytic TGFβ signalling is involved in modulating ECM structures and MMP9 is definitely one of the downstream targets of this pathway. As active TGFβ requires cleavage from its latency complex, MMP9 might play a certain role (Wilson et al., 2008). So, we performed co-immunoprecipitation which revealed a band specific to TGFβ1 expression (between 50 kDa and 75kDa) in the patients with MTLE-HS (Supplementary 1-D). This pinpointed the direct interaction of MMP9 with TGFβ1in the sclerotic hippocampi of these patients.

## Discussion

### MMP9 potentially promotes BBB damage via ZO1 degradation in MTLE-HS patient

Our findings suggested that MMP9 and its endogenous inhibitor TIMP1 were up-regulated at the transcript (mRNA) level in the hippocampus of patients with MTLE-HS. Earlier MMP9 mRNA levels were found to be up-regulated in the rat hippocampus in response to neuronal depolarisation (Rylski et al., 2009). Animal studies also suggested that increase in the mRNA expression level of TIMP1 after seizure generation was independent of de novo protein synthesis (Rivera et al., 1997). TIMP1, the endogenous inhibitor, was found to be highly up-regulated at the transcript level in the present study implying that MMP9 is tightly controlled even in patients with MTLE-HS. Therefore, the ratio of MMP9 and TIMP1 may have a role in regulating the activity of MMP9 in these patients. A prior study in children with febrile seizures and status epilepticus (SE) showed the MMP9 level in serum as well as the ratio of MMP9 to TIMP1 to be elevated (Suenaga et al., 2008). While our study showed that the protein expression level of MMP9 is significantly up-regulated in sclerotic hippocampi, the ratio of MMP9:TIMP1 greater than one indicated that MMP9 was activated in patients with MTLE-HS, further confirmed by the gelatinase activity of MMP9.

A recent report suggested mRNA level expression of MMP2, MMP3 and MMP14, TIMP 2, 3 were increased in the hippocampi of TLE patients, although mRNA expression of MMP9 or TIMP1 showed no change, and their ratio was not investigated in these patients (Broekaart et al., 2021). To the best of our knowledge, this is the first study to show the enzymatic activity of MMP9 in resected sclerotic human hippocampal tissue, while earlier reports (Mizoguchi et al., 2011, Takács et al., 2010, Li et al., 2013), showed the enzymatic activity in animal models or CSF of epilepsy patients. These results suggest that MMP9 may play a role in remodelling of ECM components leading to altered excitability. Similarly, Jourquin et al. (2003) demonstrated the increased release and activity of MMP9 in organotypic cultures when stimulated with neurotoxic kainate, while its inhibition reduced neuronal death. We found that expression of MMP9 in both glial and neuronal cells in patients with MTLE-HS, which is in contrast with animal model studies where MMP9 is expressed in dendrites and synapses of the hippocampal dentate gyrus (DG) area (Konopacki et al., 2007), and MMP9 knockout mice have less severe seizure activity (Wilczynski et al., 2008). MMP9 causes cleavage of pro-BDNF into mature BDNF in the hippocampus of PTZ-kindled wild type mice, likely inducing TrkB receptor-mediated mossy fibre sprouting, one of the pathological features of MTLE-HS (Mizoguchi et al., 2011), resulting in hyper-excitable dentate networks (Koyama et al., 2004). MMP9 inhibition reduces the density of mossy fibre sprouting by 90% (Wilczynski et al., 2008), reducing abnormal synaptic connections in epileptic animals. All of these animal studies support our findings in humans and offer a strong basis for MMP9 to be identified as a key player in the pathophysiology of patients with MTLE-HS.

Ultra-structural studies have shown that BBB abnormalities and abnormal TJs can lead to inflammation in patients with epilepsy (Cornford, 1999; Heinemann et al., 2012), whereby secreted cytokines and adhesion molecules alter endothelial cells’ TJs in a feedback manner, resulting in increased BBB permeability and damage while a lower seizure threshold ensues (Abbott et al., 2010; Librizzi et al., 2012). ZO1, which is an also a substrate of MMP9, was rapidly degraded after ischemia in wild type mice while the degradation was reduced in MMP9 knockout mice in focal ischemic model (Asahi et al., 2001). BBB permeability was increased in the cerebral hemispheres, brainstem, and cerebellum of adult rats, coupled with decreased ZO1 IR (Yorulmaz et al., 2015). Our study showed that there was loss of ZO1 IR in the hippocampi of patients with MTLE-HS. Moreover, the ZO1 expression level was found to be less in damaged endothelial lining of blood vessels in MTLE-HS as compared to intact vessels of non-seizure controls. Reduced ZO1 IR was found in kainate model after SE (Benkovic et al., 2006) while another study correlated the limit of BBB damage with duration of seizure activity, reducing the levels of ZO1 in endothelial cells (Librizzi et al., 2012). Hence, our findings are consistent with the animal studies described, demonstrating that ZO1 is one of the most critical players in maintaining the integrity of the BBB, and its loss results in barrier disruption and subsequent infiltration of inflammatory mediators into the brain. MMP9 may regulate ZO1 degradation, which in turn promotes BBB destruction.

MMP9 mediated cleavage of ECM might also influence neuronal activity and potentiate N-methyl-D-aspartate-receptor (NMDAR) currents through the activation of integrins shown in hippocampal neuronal cultures (Michaluk et al., 2009). NMDAR hyper-activation is correlated to hyper-excitability in the hippocampus of MTLE-HS patients (Banerjee et al., 2015). This hyper-activation produced by glutamate mediated by NMDARs might potentially lead to enhanced TGFβ signalling (Paul et al., 2018). Hence we delved further into the role of TGFβ signalling in MTLE-HS.

### Astrocytic TGFβ signalling might affect neuronal excitability

Earlier study had demonstrated enhanced TGFβ signalling in patients with MTLE-HS (Paul et al., 2018), where the protein expression level of TGFβ1 along with downstream effectors (SMAD3/pSMAD3) were reported. In present study, the co-expression of SMAD3 and pSMAD3 with GFAP suggested the activation of astrocytic TGFβ signalling in patients with MTLE-HS. Previous studies on animal models highlighted the association of astrocytes and TGFβ signalling focused on uptake of albumin into glial cells and binding to the TGFβ receptors present in the astrocytes following BBB opening and consequently developing seizures (Ivens et al., 2007). Also, direct application of TGFβ1 to the neocortex resulted in transcription mediated changes in astrocytic genes (David et al., 2009). TGFβ1 mediated BBB opening resulted in downward regulation of two astrocytic genes, the inward rectifier potassium channels (Kir4.1) and Aquaporin 4 (AQP4) (Perillan et al., 2002; David et al., 2009) in perivascular astrocytic end feet, associated with the BBB as part of the neurovascular unit (NVU). AQP4 knockout mice showed uncoupling of AQP4 to Kir4.1 promoting reduction in buffering of [K+] ° and prolonged seizure duration (Aronica et al., 2000; Srohschein et al., 2011) and also NMDAR mediated hyper-excitability and consequent epileptiform activity (Ivens et al., 2007). Increased TGFβ signalling also resulted in down-regulation of the glutamate transporter, Glt-1, and connexions Cx30, Cx43, and Cx26, indicating inadequate potassium and glutamate buffering (Cacheaux et al., 2009), as well as a link between enhanced excitatory synaptogenesis driven by astrocytic TGFβ signalling and BBB damage (Weissberg et al., 2015). Taking all of these studies into account, our findings imply that astrocytic TGFβ signalling may have a role in epileptogenesis in patients with MTLE-HS, possibly through BBB damage and altered neuronal functioning which needs further evaluation in patients.

Previous studies might postulate that both MMP9 and TGFβ1 function in a tandem manner and might contribute to hyper-excitability (Cacheaux et al. 2009, Heinemann et al. 2012; Kim et al. 2017). An inhibitor of MMP9, doxycycline hyclate, has been shown to prevent the loss of perineuronal nets (PNNs) after kindling in rats and delay seizure onset (Pollock et al., 2014). TGFβ1 modulates astrocytic response to pro-inflammatory mediators (Hamby et al., 2008), also regulating astrogliosis via up-regulation of ECM molecules (Smith et al., 2005). Consequently, albumin extravasation and TGFβ signalling activation in astrocytes leads to degradation of PNNs and consequent aberrant excitatory synaptogenesis (Kim et al., 2017), followed by an increase in expression of various ECM molecules like MMP9, MMP14, TIMP1, STAT3 in the hippocampus, while losartan obliterates changes in ECM via suppression of TGFβ signalling (Kim et al., 2017). Hence, MMP9 and TGFβ might interact to modulate ECM structures as evidenced by animal studies. These observations support our preliminary finding of the interaction of MMP9 and TGFβ1 in patients with MTLE-HS through protein-protein interaction. However similar interaction studies need to be further evaluated in hippocampal cultures from these patients with MTLE-HS which is beyond the scope of this study (Supplementary Fig 1).

This human study has a few limitations. One of the limitations is the small sample size of this study, which does not include age and gender-matched cases and controls. As MTLE-HS is broadly a distributed network disorder, the possibility of altered MMP9 activity and TGFβ signalling in other brain regions cannot be ruled out. Further, MMP9 activity is a dynamically regulated process and is influenced by a variety of factors. We selected the hippocampal tissues from an autopsy as well as patients for the study to reduce interference factors. Future studies addressing these limitations, particularly in animal models for TLE, will be required to validate these findings and heighten their therapeutic potential.

Summarising in the present study, we explored the potential role of MMP9 in the pathophysiology of MTLE-HS patients, specifically how it can disrupt the BBB and modulate the ECM, causing abnormal connections and fuelling epileptogenesis. Another finding of this study is that exploring the TGFβ-MMP9 interaction in MTLE-HS patients may have a synergistic effect, possibly leading to epileptogenesis. Astrocytic TGFβ signalling may have an important role in modulating the ionic distribution and leading to hyper-excitability. However, the mechanism of MMP9 activity on ZO1 is one of the many possible causes of BBB damage that has to be investigated further. Both TGFβ1 and its downstream targets modulate the ECM, and the interaction between them might lead to the possibility of MMP9 activating TGFβ1 from its latency complex in the ECM and co-ordinating in a feedback loop (Cacheaux et al. 2009; Kim et al. 2017). Remarkably, these molecules, MMP9 and TGFβ1, might prove to be better biomarkers for the prognosis of MTLE-HS. To conclude, our study revealed the novel contribution of the MMP9/TGFβ axis in the epileptogenic process of MTLE-HS patients (Figure 8). Our study not only helps us get a better understanding of the process of epileptogenesis in patients with MTLE-HS, but is also useful for finding promising drug targets and developing better therapeutic strategies.

**Figure 8:**
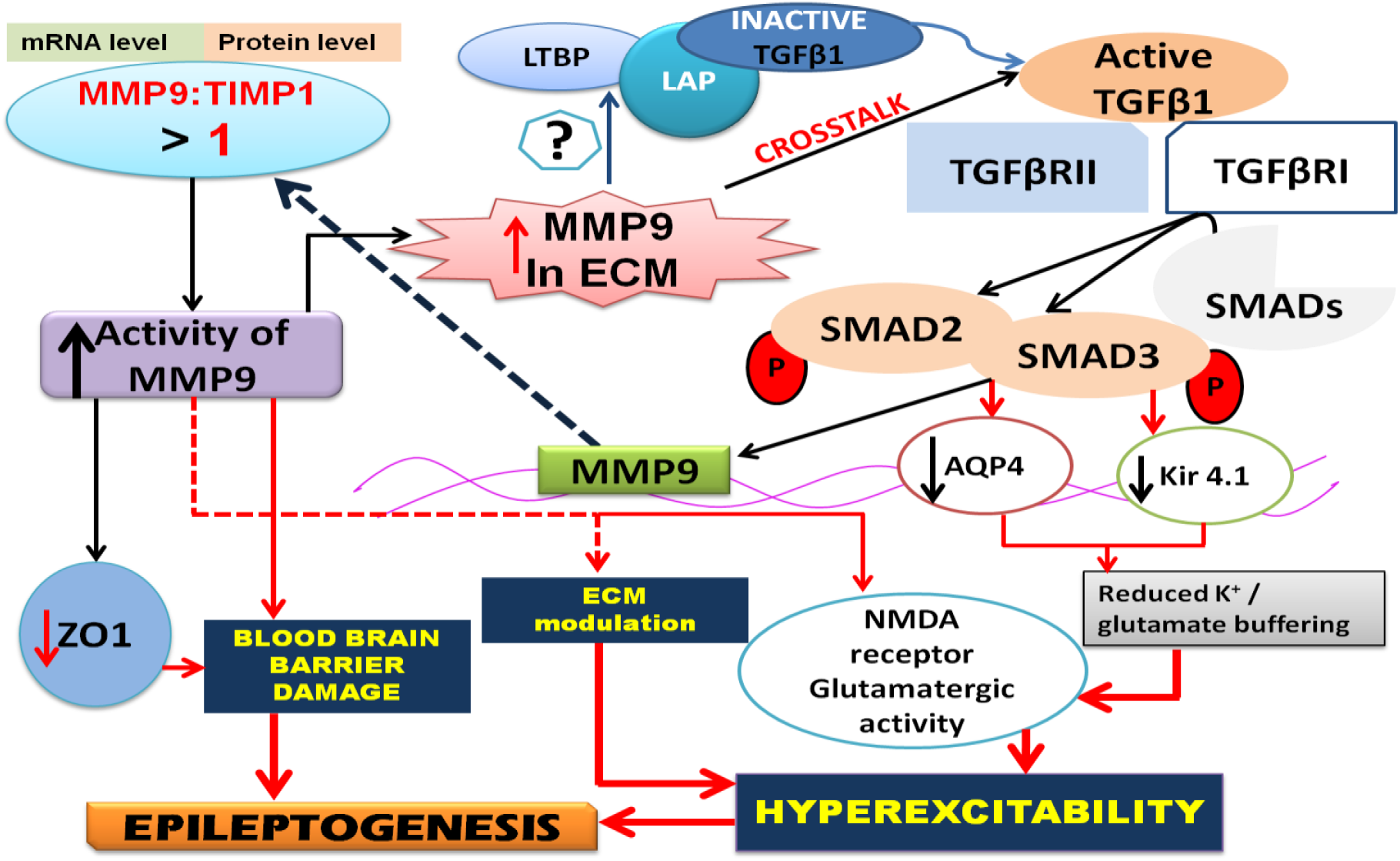
Astrocytic TGFβ signalling and high MMP9/TIMP1 ratio and their interaction promotes epileptogenesis through various processes in patients with MTLE-HS. MMP9 and its endogenous inhibitor TIMP1 alteration at transcript level was projected in patients with MTLE-HS. Also the altered protein expression of MMP9 and the ratio of MMP9 and TIMP1 have been greater than one in patients with MTLE-HS project the activation of MMP9 in these patients. The high gelatinase activity of MMP9 determined the activation of MMP9 from its pro-enzyme to its active enzymatic form which leads to increase in the active MMP9 in ECM and modulation of various ECM structures like PNNs and lateral diffusion of NMDAR contributing to hyper-excitation. Also MMP9 leads to BBB damage through cleavage of type IV collagen of basement membrane of cerebral epithelium. Low ZO1 expression, an important TJ protein of BBB and a target of MMP9, in patients with MTLE-HS suggest that MMP9 cause BBB damage through degradation of ZO1 and this loss of barrier integrity leads to neuro-inflammation. So MMP9 contributes to epileptogenesis through BBB damage, ECM modulation and hyper-excitation. On the other hand, canonical TGFβ signalling is altered in patients with MTLE-HS. Also the downstream effector molecules, SMAD3 and pSMAD3 are co-expressed with the astrocytes in patients with MTLE-HS causing alterations in glio-transmission through modifying AQP4 and Kir 4.1 channels, leading to reduced K^+^ and glutamate buffering. This might led to altered glutamatergic activity and consequent hyper-excitation and seizure generation culminating in epileptogenesis. MMP9 and TGFβ1 crosstalk in patients with MTLE-HS and MMP9, being one of the downstream targets of TGFβ signalling, might led to activation of TGFβ1 from it latency complex in the ECM. Inactive TGFβ1 is associated with LAP through LTBP which anchors the whole complex into ECM. Higher MMP9 activity in ECM may cleave this latency complex to reveal active TGFβ1 thereby promoting activation of TGFβ signalling. Hence this interaction of TGFβ1 and MMP9 in a feedback loop in patients with MTLE-HS ultimately contributes to epileptogenesis (Szklarczyk et al., 2002, Tian et al., 2007, Cacheaux et al., 2009, David et al., 2009, Michaluk et al., 2009, McRae et al., 2012, Heinemann et al., 2012, Weissberg et al., 2015, Rempe et al., 2016, Paul et al., 2018).

## Ethical Approval and Consent to participate

This study was approved by the Institutional Ethics Committee (IECPG/-40/27.11.2015, RT-3/30.12.2015) and followed the tenants of Declaration of Helsinki. An informed written consent was obtained from all the participants of the study.

## Consent for publication

All the authors have consented for publishing this article with the journal.

## Availability of data and material

The data and material is available on request to the corresponding author.

## Author’s contribution

Debasmita Paul designed the research, performed research, analysed the data and wrote the paper. Arpna Srivastava designed the research, analysed the data, and wrote the paper. Jyotirmoy Banerjee designed the research, analysed the data and wrote the paper. Manjari Tripathi designed the research and wrote the paper. Ramesh Doddamani provided clinical samples. Sanjeev Lalwani provided the autopsy samples. Fouzia Siraj and Meher Chand Sharma performed the histopathological examinations. Poodipedi Sarat Chandra provided clinical samples, designed the research and wrote the paper. Aparna Banerjee Dixit designed the research, analysed the data and wrote the paper. All authors read and approved the final manuscript.

## Acknowledgements

We would like to thank all the participants and their family for their involvement in the studies. We would also thank our technician, Md. Arman for helping with the paraffin embedded tissue block preparation and sectioning for IHC and IF.

