## Supplementary figures for "Exploring the Role of MMP9 and its Association with TGFβ Signalling in Patients with Mesial Temporal Lobe Epilepsy-Hippocampal Sclerosis"

Supplementary Figure 1: MMP9 and ZO1 expression in microvessels in hippocampal

regions;  interaction  between  TGFβ1  and  MMP9  by  co-immunoprecipitation  assay in

patients with MTLE-HS

MMP9 (IR) was observed in the endothelial cell lining of the micro blood vessels present

throughout  the  hippocampus  of  patients  with  MTLE-HS  (black  arrows)  which  indicated

expression of MMP9 in cerebral vasculature. (B) ZO1 expression was observed in intact blood

capillaries  in  hippocampal  region  of  Control  (red  arrow)  while  (C)  ZO1  expression  was

disrupted  in  damaged  blood  capillaries  in  the  sclerotic  hippocampal  area  of  patients  with

MTLE-HS (blue arrows). Scale bar 20μm. D) Co-immunoprecipitation revealed the interaction

between MMP9 and TGFβ1. Representative blot depicting the interaction between MMP9 and

TGFβ1 in the hippocampi of non-seizure controls (n=10) and the hippocampi of patients with

MTLE-HS (n=10). The MMP9 protein was pulled down through immuno-precipitation from

total protein lysate of the hippocampi of non-seizure controls and patients with MTLE-HS and

then probed for the protein expression of TGFβ1 through immunoblotting. The IP protein lysate

lane detected TGFβ1 by a band between 50-75kDa and the expression is qualitatively increased

in  patients  with MTLE-HS  which  confirms  the  interaction  of  MMP9  and  TGFβ1  in  these

patients.
